# Volumetric bioluminescence imaging of cellular dynamics with deep learning based light-field reconstruction

**DOI:** 10.1101/2022.05.31.494105

**Authors:** Luis Felipe Morales-Curiel, Gustavo Castro-Olvera, Adriana Gonzalez, Lynn Lin, Malak El-Quessny, Montserrat Porta-de-la-Riva, Jacqueline Severino, Laura Battle, Diego Ramallo, Verena Ruprecht, Pablo Loza-Alvarez, Michael Krieg

## Abstract

The application of genetically encoded fluorophores for microscopy has afforded one of the biggest revolutions in the biosciences. Bioluminescence microscopy is an appealing alternative to fluorescence microscopy, because it does not depend on external illumination, and consequently does neither produce spurious background autofluorescence, nor perturb intrinsically photosensitive processes in living cells and animals. The low quantum yield of known luciferases, however, limit the acquisition of high signal-noise images of fast biological dynamics. To increase the versatility of bioluminescence microscopy, we present an improved low-light microscope in combination with deep learning methods to increase the signal to noise ratio in extremely photon-starved samples at millisecond exposures for timelapse and volumetric imaging. We apply our method to image subcellular dynamics in mouse embryonic stem cells, the epithelial morphology during zebrafish development, and DAF-16 FoxO transcription factor shuttling from the cytoplasm to the nucleus under external stress. Finally, we concatenate neural networks for denoising and light-field deconvolution to resolve intracellular calcium dynamics in three dimensions of freely moving *Caenorhabditis elegans* with millisecond exposure times. This technology is cost-effective and has the potential to replace standard optical microscopy where external illumination is prohibitive.

## Main

Fluorescence microscopy has enabled unprecedented discoveries and became the major imaging modality in molecular and cellular bioscience. However, significant autofluorescence of the native tissues^1^ often obscures and blurs the signal from specific labels in biological samples^2^, while intrinsic photosensitivity of cells and animals, such as *Caenorhabditis elegans* ^3^, planaria ^4^ or mouse preimplantation embryos ^5^, interferes with imaging experiments that require an excitation light source. In addition, the excitation of a fluorescent protein (e.g. GFP, GCaMP) is often incompatible with an experimental design, eg. the simultaneous emission of a cyan FP (em 470nm), or if the absorption spectrum of the chromophore overlaps with that of a photosensitizers (e.g. 470nm for Channelrhodopsin^6^ and Tulips^7^). Further, the high excitation intensities that are necessary to obtain fluorescent images with extreme photon-starved samples render fluorescence microscopy potentially phototoxic and limit the lifetime of the fluorescent probe^8^. These drawbacks can be overcome by implementing bioluminescent probes as contrast labels, that can be genetically encoded to tag any protein of interest, and do not need an external excitation light source. However, traditional bioluminescent probes have a slow catalytic turnover^9^ and require a chemical cofactor, e.g. luciferin, as a photon source, which becomes oxidized prior to photon emission. More efficient luciferases based on deep sea shrimp and termed Nanolanterns, have been engineered^10, 11^. In these enzymes, the luciferase moiety is fused to a fluorescent protein, which increases the quantum yield and bears the potential to select the emission wavelength. Thus, a whole spectrum of light-emitting proteins can be tailored to a specific need. However, they require chemicals with a poor bioavailability due to low solubility in water, which greatly limits the quantum yield and concomitant signal-to-noise ratio (SNR). Thus, to obtain high-SNR images calls for long exposure times in the seconds or even tens of seconds scale which is incompatible with fast biological dynamics. Because bioluminescence imaging is widely applied for drug screening and cancer research^12^, long exposure times greatly limit the throughput.

Deep learning-based neural networks have the potential to transform microscopy research and have aided the design of advanced optical lenses^13^, autofocus ^14^, superresolution^15^, denoising^16^ and speed up complex deconvolution procedures^17, 18^ and postprocessing pipelines ^19, 20^. Here, we overcome several significant challenges, and demonstrate the use of bioluminescence as an imaging modality in the millisecond range. We constructed a new microscope with an shortened, optimized optical path, light field detection and single photo resolution in combination with machine learning that takes advantage of the development of novel cofactor chemistry^21, 22^ and transgenic animals. Because an accurate inference from a neural network, high-quality training data of biological samples is needed, we built training and inference pipelines deep learning models using two concatenated neural networks with the aim to increase the signal-noise ratio and reconstruct four dimensional information from a time series of 2D images. Despite the apparent complexity of our approach, the individual components are easy to construct, commercially available and pretrained neural networks are at everyone’s disposal. We demonstrate this approach to image nuclear dynamics in mouse embryonic stem cells, 3D imaging of zebrafish epithelial tissues and whole-body calcium imaging in muscles of free moving *C. elegans*.

## Results

Due to the low quantum yield of luciferases, standard optical microscopes are not suitable to produce bioluminescent images and dedicated instruments are commonly used^23^. Indeed, we were not able to observe any signal from Nanolanterns transfected into HeLa cells on a commercial compound microscope with the maximum exposure time of an sCMOS camera (not shown). To increase the photon collection efficiency, we thus conceived a microscope with an ultra-compact optical axis, and with a single photon-resolving qCMOS camera (Fig. 1a). With this new setup, we were able to obtain high SNR images for cells transfected with Nanolantern fusions to clathrin, actin and the plasma membrane marker lyn^11^ and supplemented with the cofactor Hikarazine (see Methods,^21^) for exposure times down to 2s (Fig. 1b, top row). Importantly, even without any further treatment, these images were comparable to fluorescence images acquired at a similar exposure time at higher magnification (Fig. 1b, bottom row) on a conventional, epifluorescence microscope. As expected, no autofluorescence was observed in the luminescence images due to the absence of external excitation light source, in contrast to the fluorescence images (Fig. 1b). We next established that the optimized bioluminescence imaging protocol enhances the photon collection efficiency in living animals. We created a transgenic *C. elegans* line that expresses a turquoise Nanolantern in their body wall muscles^24^ and immobilized individual animals for imaging on a agar pad in presence of the luciferin. In agreement with our results from tissue culture cells, we observed a strong specific signal at longer exposure time and even for exposure times down to 50 ms (Fig. 1c). Taken together, these technical improvements dramatically augmented the quantity of photons detected allowing us to significantly reduce the exposure time without additional postprocessing. Capturing the ability to record ultra-photon starved samples, we refer to our setup as ‘LowLiteScope’.

**Fig. 1:**
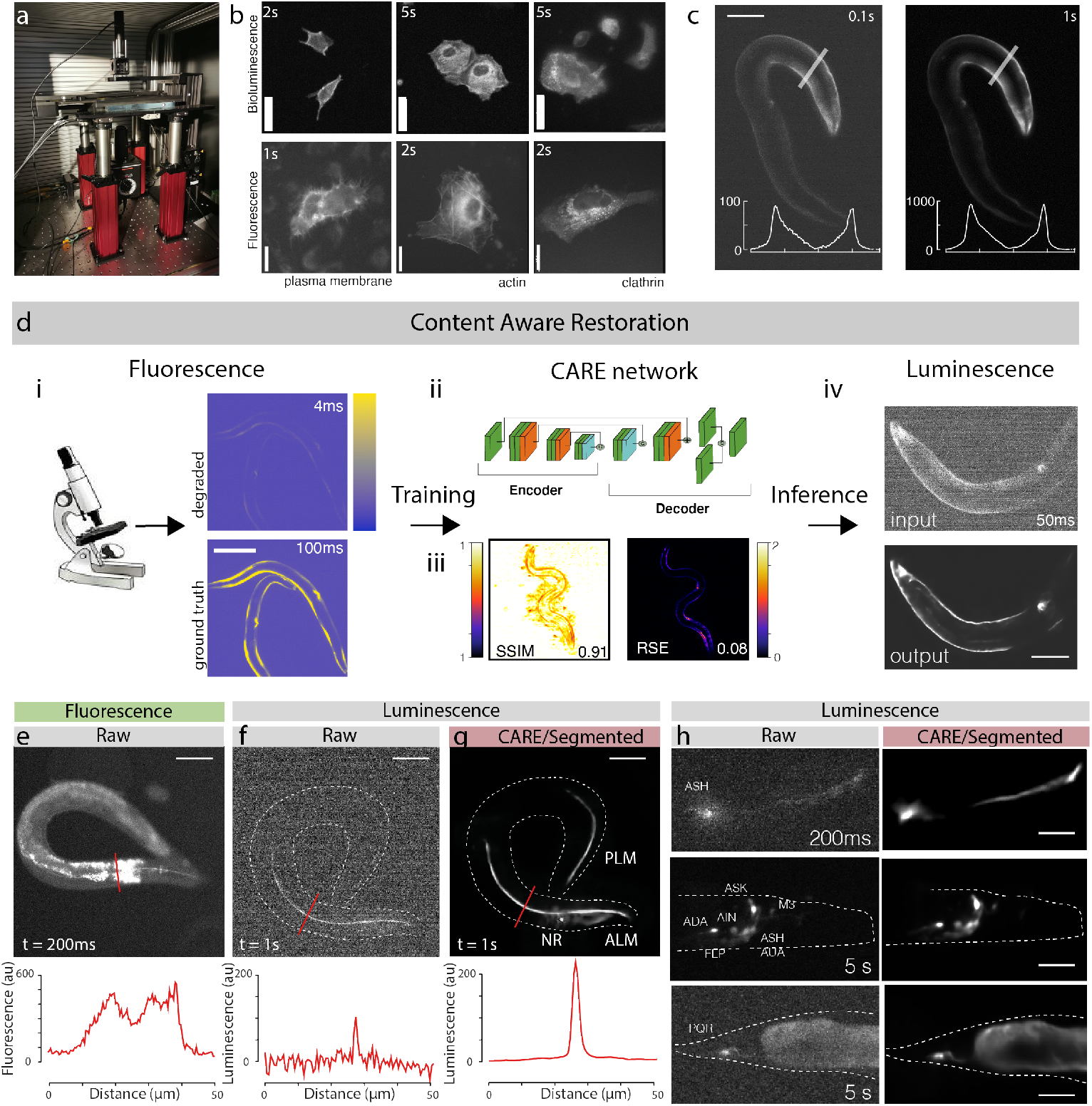
Optimized bioluminescence microscopy. **a**, Photograph of the optimized Low-Light MicroScope. **b**, Bioluminescent (top) and fluorescent (bottom) images of cell expressing the indicated marker taken on the LowLiteScope or a commercial epifluorescence microscope, respectively. Exposure times indicated in the top left of each image.Scalebar=20*μ*m. **c**, Bioluminescent images of an immobilized worm expressing a turquoise enhanced Nanolantern (TeNL) in the body wall muscle at different exposure times. Inset show the intensity profile across the line indicated in the image. Scalebar=50*μ*m. **d**, Schematic of the content aware restoration deep learning pipeline. i, pairs of images were collected in an epifluorescent microscope at different exposure times to create a low and high SNR training dataset. Note, the network was trained with pictures of animal in different body postures to avoid overfitting and memorization. ii, After subpixel registration, a deep neural network is trained to restore the test image from the high SNR ground truth. iii) Structural similarity (SSIM) and root squared error (RSE) of the predicted images vs the ground truth (see also Fig. S1). iv, The trained network is then used to restore low-SNR bioluminescent images. Scalebar=50*μ*m.**e-g**, Suppression of autofluorescence in bioluminescent restoration microscopy. Lower panel shows intensity profile through the lines indicated in the upper micrographs. e) Fluorescence picture of a worm expression mNeonGreen-enhanced Nanolantern (GeNL) in touch receptor neurons; the same transgenic in bioluminescent contrast f) before and g) after AI-denoising. Scalebar=50*μ*m. **h**, Versatility of the neuronal reconstruction as shown on several neurons in *C. elegans*, such as ASH, a neuronal ensemble expressing the Turquoise-enhanced Nanolantern in glutamatergic neurons (*eat-4*p:TeNL) and PQR. Scalebar=15-30*μ*m.

Inspired by these positive results, we set out to test the advantages of low-background autofluorescence recordings, and established transgenic *C. elegans* animals that express a green enhanced Nanolantern exclusively in the touch receptor neurons (TRNs). High resolution imaging of these neurons is often precluded by the abundant autofluorescence that emanates from the ubiquitous gut granules under epifluorescent illumination (Fig. 1e). This is of particular concern in old animals^25^, in which the signal of the autofluorescence can become more intense than the specific label, which makes it difficult to distinguish between both. Indeed, the fluorescent images derived from animals expressing GFP in TRNs on a standard epifluorescence microscope showed extensive out of focus light due to background autofluorescence of the gut (Fig. 1e). In contrast, living animals that express the TRN::luciferase and were supplied with the optimized cofactor, a single, specific signal is visible from the monopolar dendrites of these neurons, and no spurious autofluorescence can be seen (Fig. 1f). However, because of the small size of the TRNs, the obtained SNR is very low at exposure times as short as 1s. Because there is no noise inherent to the sample (only from the detector), we reasoned that prior knowledge of the underlying sample structure should facilitate superior image reconstruction with dramatically improved SNR using deep learning-based content aware image restoration (CARE) algorithms^16^. To clean up these images and increase the effective SNR, we combined these bioluminescent microscopy images with convolutional neural network that transforms a degraded image to a desired high quality target given a proper training with a known signal distribution^16^.

Because CARE models for *C. elegans* and bioluminescence are in-existent, we first developed a generalizable training pipeline to predict high quality, ground truth images from noisy input. We thus collected image pairs derived from fluorescence microscopy acquired at extremely low exposure times reflecting the noisy input with poor SNR and used high SNR images from long exposure times as the ground truth target (Fig. 1d i). The training data set consisted of animals transgenic for mTurquoise body wall muscles, which showed high specific signal and lack of any visible autofluorescence within the region of interest. To achieve a high variety of body postures and thus 2D intensity distributions, we recorded the fluorescence signal from the body wall muscles in freely moving animals. After data collection and preprocessing, we varied the hyperparameters to find the optimal network configuration^16, 26^ specific for our training dataset (Fig. S1, 1d ii), and evaluated its out-of-sample performance on unseen noisy images using the structural similarity (SSIM) and residual squared errors (RSE) metrics for training quality (Fig. 1d iii, Supplementary Fig. S1 and Methods). We then built an inference pipeline with the model showing the highest confidence and lowest error to predict the ground truth from the noisy bioluminescent images which turned out to be completely clean and devoid of artifacts (Fig. 1d iv). When we applied this model to degraded bioluminescent images derived from transgenic TRNs in aged animals, our model was able to effectively enhance the SNR and cleanly visualize the neurons for further inspection (Fig. 1g). Importantly, this approach is not limited to a specific neuron in *C. elegans*, and we have successfully enhanced the degraded bioluminescent images acquired for ASH, PQR and vGLUT EAT-4 expressing neurons in the head (Fig. 1h, ^24, 27^). Strikingly, we found satisfactory performance of the model with exposure times as low as 200 ms taken on ASH, a neuron in the head of *C. elegans* with a diffraction-limited axon caliper. Taken together, the combination of optimized optical path, cofactors^21, 28^ and dedicated machine learning algorithms from the CARE family enabled the acquisition of high-SNR images at exposure times as low as 50 ms in living animals and tissue culture. Importantly, we showed that we could achieve high-performance with a small, but diversified training dataset, that resulted in a generalizable and transferrable model to infer noiseless images from severely degraded inputs of different cellular structures in *C. elegans*. Consequently, this allowed us to build our pipelines using free cloud-computing resources, which are accessible to a standard research laboratory^26^.

Photo-bleaching during fluorescence microscopy is an indicator for potential photo damage to the cell^8^, especially at lower wavelength commonly used for one photon live-cell imaging. Without the requirement of an excitation light source, bioluminescence has the advantage to circumvent potentially phototoxic effects^29^. We thus compared the activation of a cellular stress reporter in fluorescence and bioluminescence microscopy and generated transgenic animals expressing mNeonGreen-NanoLantern fused to DAF-16, a promotor of longevity^30^ and reporter for various stresses, including reactive oxygen species^31^. To verify that the bioluminescent stress reporter signals the animal’s exposure to cytotoxic stresses, we followed cytoplasmic/nuclear shuttling during the application of a heat shock and compared it to the stress response after fluorescence imaging. With both reporters, we observed a strong nuclear relocalization immediately after the heat shock (Fig. 2a,b; Video S1). Even though we only detected muscle and neuronal cells in the bioluminescent animals, the activity of the bioluminescent reporter was more pronounced. We reasoned that the absence of autofluorescent background in the bioluminescent images enabled a higher dynamic range. When we omitted the heat shock, the unstressed, control animals that were recorded with fluorescence microscopy showed a slight but significant increase in nuclear DAF-16 localization (Fig. 2b,c; R=0.95, p*<*1e-15), which was strongly reduced in the bioluminescent images (Fig. 2c; R=0.56, p=1e-5). We thus speculated that the spontaneous activity if DAF-16 might be triggered by the reactive oxygen species during the fluorescence illumination^8, 32^. Taken together, the application of bioluminescent reporter offers a higher dynamic range in absence of background autofluorescence and could possibly guide the discovery of stress pathways that would otherwise be obscured by the cellular response to external light.

**Fig. 2:**
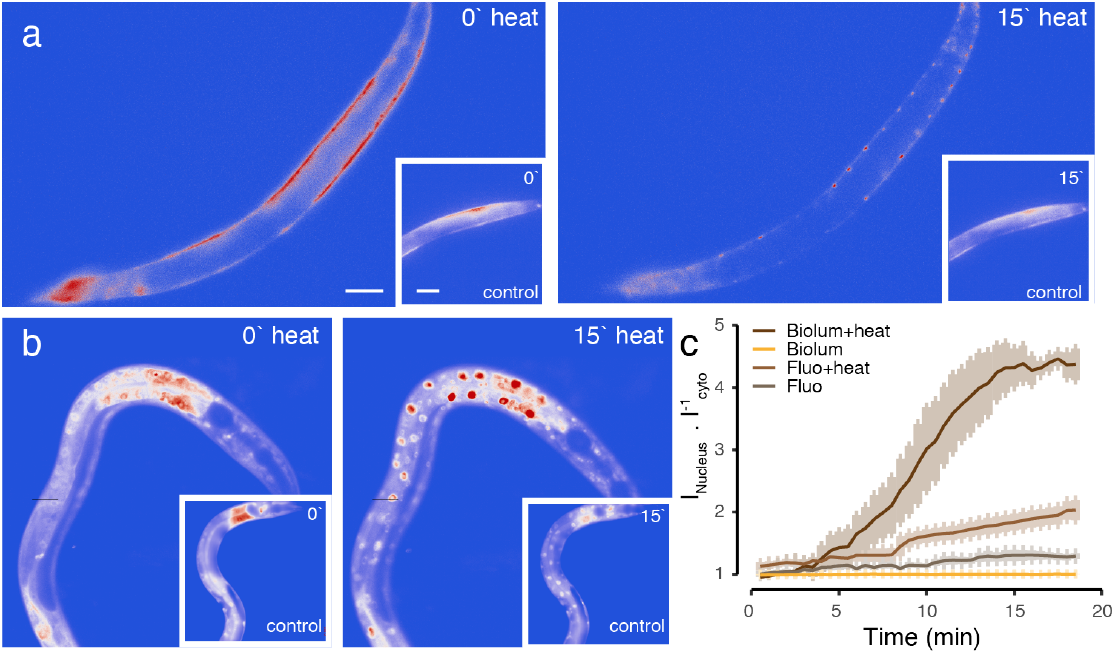
Stress reporter activation under external illumination **a**,**b** Bioluminescent (a) and fluorescent (b) DAF-16::GeNL before (0’) and after (15’) exposure to 37C heat stress. Inset shows animals at the same timepoints without external heat stress.**c** Quantification of the nuclear/cytoplasmic DAF-16 ratio over 18 min of the experiment in the four tested conditions. Scale bars = 40μm.

To demonstrate that our combination of bioluminescent imaging and deep learning can be generalized to other animals and biosystems, we generated bioluminescent zebrafish embryos expressing a membrane-bound red shifted nanolantern and mounted them for imaging in our LowLiteScope at 4 hours post fertilization. Under fluorescence excitation, the EVL is clearly visible as a tessellated epithelial cell layer (Fig. 3a). Under bioluminescence contrast, however, strong out-of-focus haze limited the signal strength and SNR, even though we were able to record a signal reminiscent of the cell junctions after 100ms exposure time (Fig. 3b, Supplementary Fig. S2). We thus combined the final images with a pre-trained CARE model, originally established for epithelial monolayers in *Drosophila* wing discs^16^, a tissue with similar tessellated morphology and signal distribution. Despite the challenging task due to the poor input SNR, we found that this model was generalizable and fit our input extremely well, being able to greatly improve the SNR (Fig. 3b,c). Critically, these signal restorations and improvements enabled the segmention of individual cells in the embryo (Fig. 3c) which afforded the calculation of their perimeter and cell area (not shown) - a procedure that otherwise would not have been possible.

**Fig. 3:**
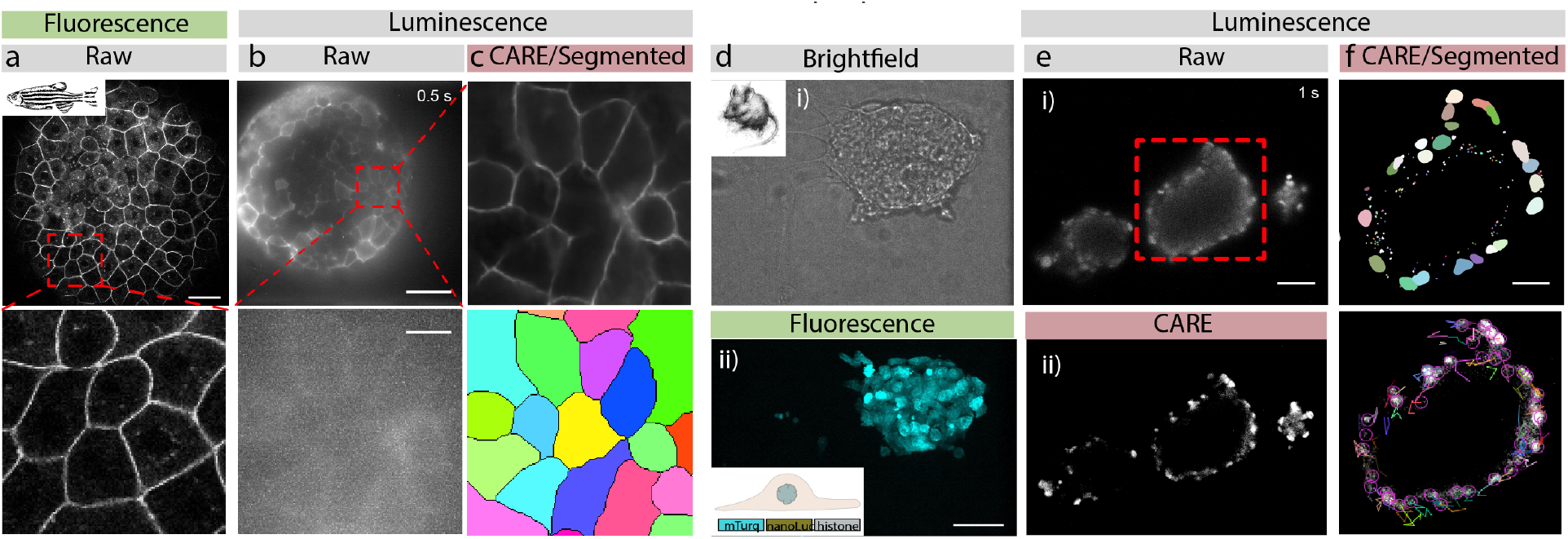
Seamless denoising and segmentation of bioluminescent samples **a**, Laser scanning confocal fluorescence image of a 4hpf embryo expressing membrane bound GPI-GFP. The red box indicates the close-up below.**b**, Unprocessed bioluminescence image of a 4hpf zebrafish embryo expressing GPI:GFP targeted to the plasma membrane. The red square indicates the high magnification close up below. **c**, The same bioluminescent signal of the embryo was restored using a pre-trained CARE pipeline optimized for epithelial monolayers. The bottom picture shows the segmented bioluminescent image. No segmentation was possible on the raw image. **d**, Brightfield (top) and fluorescence image (bottom) of a spheroid of mouse embryonic stem cells. **e**, Unprocessed raw (top) and denoised (bottom) bioluminescence image of similar spheroids. **f**, Segmented nuclei (top) after denoising, overlayed with their individual tracks throughout the timelapse (bottom).

We were next interested to demonstrate subcellular dynamics in mouse embryonic stem cells and generated a stably transgenic cell line expressing a nuclear localized luciferase by fusing mTurquoise-NL to histone (see Methods, Fig. 3d). After optimization of the co-factor delivery (see Methods) we performed timelapse imaging of individual cells in spheroids from mESCs and recorded their nuclear dynamics (Fig. 3e, Video S2). Expectedly, the images were noisy, due to the low quantum yield. To improve visual quality and the ability to quantify nuclear trajectories, we sequentially employed two published convolutional neural networks. We first passed the noisy images through a pretrained CARE neural network for denoising nuclear morphologies^16^ and then performed nuclear segmentation with the StarDist algorithm ^19^ (Fig. 3e, f). This approach allowed us to track the migratory path for each individual nuclei within bioluminescent spheroids. Taken together, these approaches demonstrate the possibility to image subsecond dynamics of subcellular localized bioluminescent probes in *C. elegans*, zebrafish and mouse embryonic stem cells.

Up to this point, the long exposure times in bioluminescence imaging have largely hindered the acquisition of three dimensional image stacks, especially in moving animals. Often, it is desirable or even important to obtain the whole 3D representation of a fast biological process, e.g. during calcium imaging of neuron or muscle contraction. We thus sought to establish single-exposure volumetric light field imaging^33^ to quantify calcium dynamics in freely moving animals using bioluminescent calcium indicators. To do so, we equipped our LowLiteScope with a microlens array that is matched to the magnification and numerical aperture of the imaging lens and projected the entire light field onto the qCMOS sensor for plenoptic imaging in three dimensions (Fig. 4a). To obtain the 3D information from a 2D image, the light field needs to be deconvolved computationally^34, 35^. Traditionally, however, this process is computationally very demanding and takes up to several minutes for a single image^36^ amounting to many hours or even days computing time for a whole time series, which makes recording of cellular dynamics unattainable. Several AI-based algorithms have been proposed to speed up the deconvolution and enhance performance^18, 37, 38^, that significantly outperform traditional light field processing. To create a neural network for the reconstruction of *C. elegans* expressing a fluorescent calcium reporter in the body wall muscles, we first trained a NN with synthetic light field data^37^ as the input and experimental confocal stacks as the target (Fig. 4b). Both features were derived from the same immobilized animal, each stack was convolved with the light field point-spread-function (PSF) to generate the synthetics input data (Fig. 4b and Methods). We then extended the model using transfer learning with purely experimental data containing fluorescent light field images and *z*-stacks taken from the same sample as just described. This network knowledge expansion allowed us to be more specific to the experimental images we got from our setup, gave more flexibility to perform well for low and high SNR light field images, reduced the possibility to obtain artifacts and improved the inference quality (Fig. 4 c). As described before^37^, this approach shortened the processing time from 30h to 100ms per full frame image as compared to traditional lightfield deconvolution algorithms. We then used this model to reconstruct the bioluminescent light field images (Fig. 4d). We found that exposure times of 5s are required to obtain a clear representation of the scene, but with blurred dynamics due to sample movement. In order to enable faster frame rates to ‘freeze’ animal movement and capture the full dynamics of the calcium dye, we applied the CARE pipeline for denoising the low SNR lightfield images obtained at low exposure times prior to the light field deconvolution within a sequential application of two neural networks (Fig. 4e). The denoising led to a striking increase in SNR of the light field image, and with this approach, we were able to obtain significantly better reconstructions than without (Supplementary Fig. 5). Moreover, we were able to obtain three dimensional calcium recordings from whole animals with typically 200-500ms exposure time to create a full 3D stack of the bioluminescent scene which corresponded to a z-resolution of 1.5 um and reconstructed 31 z-planes in conventional widefield microscopy (Fig. 4f, Supplementary Fig. S5). Strikingly, this is equivalent to 6.4 ms exposure per frame in traditional volumetric imaging. In these bioluminescent calcium recordings, we observed higher intensity on the concave side of the bend, consistent with high-calcium concentration during muscle contraction (Video S3). Importantly, the reconstructions preserved the relative intensity distribution within the sample, as we did not find significant differences between the reconstructed forward projection and the ground truth (Supplementary Fig. S3). We also observed that most calcium signal comes from equatorial region of the muscles and rapidly drops off towards the lateral sides (Fig. 4g). This implies that the contractile power is localized to the equatorial regions, where the largest bending moment can be applied. Consistent with a high calcium concentration during muscle contraction, we observed that the largest intensities mapped to positive body curvatures (Fig. 4h).

**Fig. 4:**
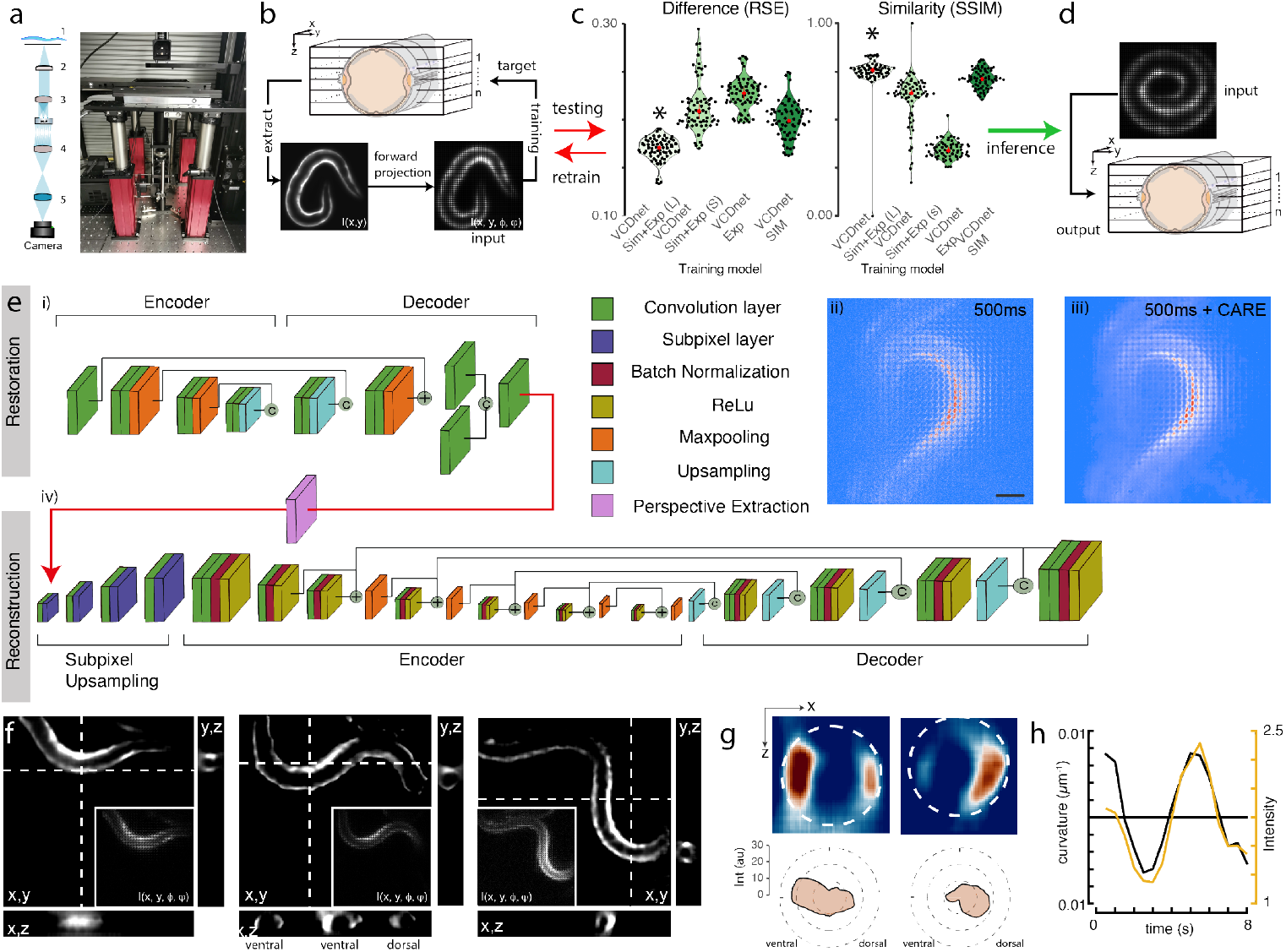
Single-exposure, volumetric bioluminescence microscopy **a**, Schematic and photograph of the optimized Low-LightField MicroScope. 1) Sample, 2) Objective, 3) Tube lens, 4) Microlens array, 5) Relay lens. **b-d**, Training pipeline to obtain fast deconvolution of 2D experimental lightfield data into 3D image stacks. b) A 3D image stack was acquired on fluorescent samples representative of the bioluminescent signal in the final experiment. The stack was convolved with the lightfield PSF, to obtain a synthetic lightfield image, which was subsequently used to map onto the 3D ground-truth stack. The training quality of the individual models (c) was tested against unseen samples by calculating the difference and similarity to the ground truth. d) The best model with the lowest error and highest similarity was used to reconstruct experimental bioluminescence images. **e**, Pipeline for bioluminescence reconstruction is composed. An initial i) CARE denoising step is used to increase the SNR of ii) noisy bioluminescent lightfield images. The individual layers are color coded according to their function. The clean images (iii) are fed into the VCD network^37^ (iv) after perspective extraction to reconstruct the 3D information. Scalebar = 50*μ*m. **f**, Sequence of reconstructed 3D images of a moving animal showing high calcium activity at its contracted side. Images show a single plane of the reconstructed z-stack. Inset corresponds to the raw lightfield image. **g**, Sideview image of the curvature-dependent calcium signal in muscles during ventral and dorsal body bends. The polar plot shows the intensity distribution on the ventral and dorsal side. Dotted line corresponds to the circumference of the animal. **h**, Intensity of the bioluminescent calcium indicator and curvature variation on the ventral side during animal crawling under the lightfield microscope. Black=curvature, yellow=calcium signal.

Taken together, we have shown that a combination of an optimized optical path and advanced computational tools dramatically improves the SNR of bioluminescence microscopy that rivals that of conventional fluorescence microscopy. We have demonstrated the performance on living tissue culture cells for subcellular labeling of actin and microtubules, zebrafish epithelial cell organization and nuclear dynamics in mouse embryonic stem cells. Lastly, we combined a sequential neuronal network composed of content aware image restoration pipelines and light field reconstruction to enable high-speed, subsecond volumetric imaging of a genetically encoded calcium sensor in freely moving animals. Novel luciferases and cofactors will be needed to obtain single cell resolution at high magnification in crowded tissues, e.g. organoids or for whole brain luminescent calcium imaging. In the future, spatiotemporal resolution and light-capturing ability could further be improved through combinations of wavefront coding^39^, tunable optics^40^, Fourier-lightfield microscopy^41^, and new transformer networks that are trained to provide a spatially super-resolved representation of the scene^42^. Our results pave new avenues for excitation-free, non-invasive low light imaging in microscopy, diagnostics and biomedicine.

## Data and Code availability

All training data and bioluminescent source images will be deposited on zenodo.org upon acceptance of this article. The code of the training and inference pipelines and instructions to run it will be freely available under this link.

## Acknowledgments

We would like to thank the NMSB and SLN lab for discussions and suggestion on optical design and imaging procedures. We thank Senda Jiménez-Delgado and Neus Sanfeliu-Cerdan for help with molecular biology and Valeria Venturini for zebrafish injections. MK acknowledges financial support from the ERC (MechanoSystems, 715243), HFSP (CDA00023/2018), Spanish Ministry of Economy and Competitiveness through the Plan Nacional (PGC2018-097882-A-I00), FEDER (EQC2018-005048-P), “Severo Ochoa” program for Centres of Excellence in R&D (CEX2019-000910-S; RYC-2016-21062), from Fundació Privada Cellex, Fundació Mir-Puig, and from Generalitat de Catalunya through the CERCA and Research program (2017 SGR 1012), the Laserlab-Europe (H2020 GA no. 871124) in addition to funding through H2020 Marie Skłodowska-Curie Actions (754510 to AG, and 847517 to LFMC).

## Author contribution

LFMC, GC and MK built the microscope; LFMC, ACG, MPR, LL and MEQ performed experiments; LFMC wrote software and performed the deep learning; JS, LB, MPR performed transgenesis of mESCs and *C. elegans*; LB, VR, DR, PLA, MK supervised and MK conceived the project. LFMC and MK wrote the first draft, with input from all authors.

## Supplementary Videos

**Supplementary Video 1** Dynamics of the DAF-16 transcription factor in response to external heat. For display purposes, the video was denoised using the deep learning pipelines developed in this manuscript.

**Supplementary Video 2** Dynamics of mouse embryonic stem cells within a spheroid. For display purposes, the video was denoised using the deep learning pipelines developed in this manuscript.

**Supplementary Video 3** Three dimensional calcium dynamics of a freely moving animal. The video was denoised and reconstructed from a 2D lightfield image using the deep learning pipelines developed in this manuscript.

