## Supplementary Material for "Volumetric bioluminescence imaging of cellular dynamics with deep learning based light-field reconstruction"

### 1 Supplementary Methods

***C. elegans* culture** Animals were maintained on Nematode Growth Medium (NGM) plates seeded with *Escherichia coli* OP50 bacteria using standard protocols[43]. Where indicated, age-synchronized animals were used and handled as described[44]. The following strains have been used: MSB266[*eat-4(ky5)III; hpIs166; mirEx89[pNMSB26(sra-6p::sng-1::TeNL250::unc-54 3'UTR)*],
MSB343[*mirEx123(myo-3p::TeNL250::unc-54)* 5 ng/ul],
MSB426[*mirSi16 II; eat-4(mir28)III; mirSi15 IV; lite-1(ce314)X, mirEx168[pNMSB17(eat-4p::TeNL250)*,
MSB557*[mirEx218[pNMSB52([mec-4p::GcNL::let-858 3'UTR)*,
MSB577[*bus-17(br2)X; mirEx123(myo-3p::TeNL250::unc-54)*],
MSB1041[*bus-17(br2)X; mirIs92(daf-16p::daf-16::GeNL::unc-54)*]

**Molecular Biology** All plasmids used for this study were constructed using the Gibson assembly method. pNMSB17, pNMSB26 and pNMSB40 plasmids carrying the turquoise-enhanced Nanolantern (TeNL) have been described elsewhere[24]. To generate pNMSB52 (TRN:GeNL) a mNeonGreen enhanced NL (GeNL) gene was cloned with primers 5'-atggtctccaagggagaggaggacaac-3' and 5'-TTACGCGAGGATACGCTCGCAGAGAC-3' into a plasmid containing the *mec-4* promoter.

The membrane-bound orange enhanced Nanolantern was subcloned from Lyn-OeNL_pcDNA3 (Addgene plasmid # 89528) into a GPI-anchor containing plasmid. In vitro transcribed mRNA was injected at 100pg into one cell stage zebrafish eggs.

**Transgenic mouse embryonic stem cell line generation and maintenance** To generate the transgenic mouse embryonic stem cell (mESC) line, 1 x $10^6$ G4 cells [45] were lipofected using Lipofectamine™ 3000 Transfection Reagent (Thermo Fisher Scientific, L3000001) mixed with 0.625 $\mu$g of PX330 (Addgene, 98750) (sgRNA 5'-ACTGGAGTTGCAGATCACGA-3') and 1.875 $\mu$g of circular HDR template. Lipofection was carried out using a PenStrep-free mESC medium (described below). Cells were single-cell FACS-sorted for mTurquoise expression using a BD Influx Cell Sorter (646500KZ, model Influx V7 Sorter, software BD FACS Sortware). Individual clones were screened by PCR amplification.

mESCs were maintained and expanded on 0.2% gelatin-coated dishes in mESC media composed of the following: Knock-Out DMEM (Thermo Fisher Scientific, 10829-018) supplemented with 15%Fetal Bovine Serum (FBS) (in-house mESC-tested), 1,000 U/ml Leukemia Inhibitory Factor (LIF, in-house generated), 1 mM Sodium Pyruvate (Thermo Fisher Scientific, 11360070), 1x MEM Non-Essential Amino Acids Solution (Thermo Fisher Scientific, 11140050), 50 U/ml penicillin/streptomycin (Thermo Fisher Scientific, 15140-122) and 0.1 mM 2-mercaptoethanol (Thermo Fisher Scientific, 31350010). Cells were cultured at 37°C with 5%CO2. Medium was changed every day and cells were passaged using 0.05% Trypsin-EDTA (Thermo Fisher Scientific, 25300054) and quenched 1:5 in DMEM supplemented with 10%FBS (Life Technologies, 10270106). Cells were tested monthly for mycoplasma contamination by PCR.

**Tissue culture experiments** The luminescent markers clathrin:CeNL, actin:YeNL and the plasma membrane marker lyn::OeNL[11] were transfected into HeLa cells with Lipofectamine using standard procedure. Three days post transfection, tissue culture cells were supplemented with Hikarazine cofactor (at 0.8 $\mu$g/ml) and imaged in the LowLiteScope. CeNL-Clathrin_pEGFP, ReNL-Actin_pcDNA3 and Lyn-OeNL_pcDNA3 was a gift from Takeharu Nagai (Addgene plasmid # 89540, # 89531, # 89528).

**Zebrafish** Zebrafish (Danio rerio) were maintained as previously described [46]. Embryos were kept in E3 medium at between 25° and 31°C before experiments and staged according to morphological criteria [47] and hours post fertilization (hpf). Wild-type embryos were obtained from the AB strain background. All protocols used have been approved by the Institutional Animal Care and Use Ethic Committee (PRBB–IACUEC) and implemented according to national and European regulations. All experiments were carried out in accordance with the principles of the 3Rs (replacement, reduction, and refinement).

### Bioluminescence microscopy

#### 1.0.1 Instrument design

We redesigned a standard widefield microscope and with a minimal set of components to minimize photon loss. We used a 40x/1.25 silicon immersion lens (Olympus) as imaging objective, and a 100 mm tube lens (ASI Imaging) to project the collected light onto a single photon resolving camera (Hamamatsu Orca Quest, or Fusion). The sample was mounted on a standard K-frame on top of a motorized Maerzhaeuser stage and a tissue culture incubator for temperature and $CO_2$ control. To keep the sample in focus, an objective piezo-positioner was used (MIPOS 500, Jena Piezosystems), in conjunction with an autofocus system (CRISP, ASI Imaging). The whole setup was controlled with PycroManager[48], calling $\mu$Manager out of Python.

For single exposure volumetric imaging we added to the setup a microlens array (200x200 lenses) with a pitch of 222$\mu$m and focal length, a radius of curvature of 0.85 mm, a focal distance of 1.86 mm and a size of 11 mm $\times$ 11 mm $\times$ 1.5 mm (Okotech APO-Q-P222-F1,86 (633)). We chose these specific properties to match the NA of the described objective to ensure optimal spacing between the lenslets. To align the focal plane of the microlens array with the image plane of the telescope, we used a relay lens composed of two lenses of 100 mm. We used a zoom housing system to modify the distance between the CMOS sensor and the lenses in order to calibrate the setup before the image acquisition.

#### 1.0.2 Image acquisition

**Mouse embryonic stem cells (mESCs)** Days prior to imaging, $10^6$ transgenic mESCs were plated on a 35 mm No. 1 glass bottom $\mu$-Dish (Cellvis, D35-10-1-N) coated with 0.1% gelatin (Stem Cell Technologies, 07903). Cells were 40-85% confluent during imaging, providing variation in 3-dimensional colony (spheroid) volume and size. Cells were washed with PBS and new mESC media was added immediately prior imaging to eliminate debris. To image bioluminescence, the LowLightScope was equipped with environmental control to maintain 37°C and 5% $CO_2$ within a top stage incubator (LCI incubator system, Gas Mixer CA-10 and temperature controller TP10) and a qCMOS Orca quest C15550-20UP camera. 2-4 $\mu$M fluorofurimazine (FFz, Promega) subtrate was perfused through the cell culture dish, using a peristaltic pump, at a rate of 1 mL/min. Bioluminescence was observed immediately following substrate perfusion. For timelapse imaging, fresh substrate was added every 45-60 mins. Z-stacks of 5 $\mu$m spacing were acquired every 5 minutes with an exposure time of 500 ms per slice.

***C. elegans*** Bioluminescence images were acquired on the optimized LowLiteScope described above. If otherwise stated, L4 or young adult animals were mounted in a 1% agar pad and treated with citrate buffer (pH 6.5), 20% DMSO, 0.05% triton X-100 and 1.25% pluronic F128 and the chemical cofactor hikarazine[28] for the generation of a bioluminescence signal.

**Daf-16 in *C. elegans*** Bioluminescence Daf-16 dynamic was acquired on the optimized LowLiteScope described above. Young adult animals were mounted in a 1% agar pad and 1 $\mu$L of 20 $\mu$M fluorofurimazine (FFz, Promega) was added directly prior to imaging without preincubation time. We acquired the dynamics with and without exposing the animal to a heat shock. We used a stage-top incubator system T (LCI), to maintain a temperature of 37 °C during imaging.

Fluorescence Daf-16 dynamic was acquired on a Leica DMi8 inverted fluorescence microscope with a Hamamatsu Orca Flash4.0 V3 sCMOS and a 40x/1.1 water immersion lens (Leica). Animals were mounted in a 1% agar pad. Same as above, we acquired the dynamics with and without exposing the animal to a heat shock. We used an incubator (Warner Instruments) to maintain a temperature of 37 °C during imaging. Animals were excited at 488-nm using a Lumencor SpectraX LED illuminator guided through a triple band pass dichroic (FF459/526/596-Di01-25x36) and images were taken every 30s with a 100ms exposure time.

For final quantification, the intensity of an ROI drawn around the nucleus was divided by the the intensity of the cytoplasm close by, except for the unstressed bioluminescence animals where the nucleus was not visible. The nuclear/cytoplasmic ratio was plotted in R and displayed in Fig. 2c.

**Zebrafish** AB wild-type zebrafish embryos were micro-injected at 1-cell stage with 100 pg GPI-GFP mRNA. mRNA was synthesized using the mMessage mMachine Kit SP6 Kit (Ambion AM1340M). Embryos were dechorionated at sphere stage (4 hfp) and mounted in 1% low-melting point agarose in Danieau´s solution (58 mM NaCl, 0.7 mM KCl, 0.4 mM MgSO4, 0.6 mM Ca(NO3)2, 5 mM HEPES) on 35 mm glass bottom dish (MatTek). Embryos were imaged with a 20X glycerol-immersion objective on Leica SP8 confocal microscope. Laser excitation of

488 nm and HyD detector were used. Z-stacks of 0.2 $\mu$m spacing between z-slices were acquired.

### Machine learning

#### 1.0.3 Content aware restoration (CARE)

CARE has proved that is possible to effectively denoise fluorescence microscopy images under conditions of low light or low exposure time[16]. One source of the distortions in low SNR fluorescence images are typically limitations of the camera read-out noise, photon noise or the resolution loss due to under-sampling. Image denoising is the process of separating the signal $s$ and the signal-degrading noise $\nu$ of a distorted image $x$. The noisy image can be thought of being the result of a function $x = f(y)$ applied to the ground-truth (GT) image $y$. The inverse function, e.g. $y = f^{-1}(x)$ is typically computationally demanding or intractable to describe mathematically. Instead of calculating the real denoise function $f^{-1}$, CARE learns to map an approximation of this function by processing a large set of pairs ($x$,$y$) of noisy images $x$ and their corresponding true images $y$ by using a CNN based on residual U-net architectures [16].

**Denoising bioluminescent body wall muscle and neurons in *C. elegans*** For training data acquisition, L4 transgenic C. elegans expressing mTurquoise fused to nanolantern (MSB343) were imaged on a Leica DMi8 inverted fluorescence microscope with a Hamamatsu Orca Flash4.0 V3 sCMOS using different camera exposure times. For imaging, an epifluorescence illummination with 25x×/0.95-NA (numerical aperture) water-immersion objective was used. Fluorescent animals were excited at 430-nm using a Lumencor SpectraX LED illuminator guided through a triple band pass dichroic (FF459/526/596-Di01-25x36). To obtain a low SNR image representing the bioluminescent setup, we used a low exposure time (4 ms), and to obtain a high SNR image representing the 'ground truth' and the desired inference quality of our model, we used a high exposure time (100 ms). Both channels were acquired simultaneously through a Hamamatsu Gemini W-View optical beamsplitter equipped with a CFP/Venus emission filter set for the ground truth and degraded channel, respectively. The two images were cropped and superpositioned in python as part of the preprocessing pipeline prior to the training. Importantly, both channels were recorded at the same time allowing us to use freely moving animals as training sample. To achieve a high variety of postures, animals were mounted in a ”pool” created with 1% agarose pad and halocarbon oil 700 (SigmaAldrich) to capture different postures of the worm, recapitulating crawling movement. This method enables data set diversification, permitting the model to generalize in freely moving animals. We collected 2500 pairs of images of average size $1024 \times 512$, resulting in a mere $\approx 9$ GB of training data.

From the 2500 images, we extracted subimages (patches) which were given to a U-Net type architecture (Denoising CARE 2D topology) for training and validation. We tried different hyperparameters to optimize the configuration of the model and consequently, improve the model performance, e.g minimize the loss function. After training we evaluated the models with 100 unseen images of *C. elegans* at various positions. Please refer to Supplementary Fig. 1a for more details. To validate the quality of the prediction, we compute the RSE and SSIM against unseen GT images.

**Denoising, segmentation and tracking of mouse embryonic stem cell** To restore the bioluminescence stem cell timelapse, we use the *human U2OS cells* dataset stained with Hoechst 33342 markers and imaged with a camera exposure range of 15 to 1000 ms provided by the authors of Reference [49]. We followed [16] training protocol and inferred our embryonic stem cell data with the best model checkpoint we obtained. We saw a considerable improvement after the processing of every frame in our timelapse. Exploiting this, we further analyze processed the data by using a pretrained Stardist segmentation algorithm [19] and TrackMate tracking algorithm [50].

**Denoising, projection and segmentation of bioluminescent zebrafish embryo** For the bioluminescent Zebrafish embryo, 100 pg of lyn::OeNL mRNA was injected in 1-cell stage embryos to visualize the plasma membrane. Approximately two hours post-fertilization (2 hpf) at the 64-cell stage, embryos were incubated in 400 $\mu$M fluorofurimazine (FFz, Promega) for 2 hours. At sphere embryonic stage (4 hpf), embryos were mounted in a 35 mm No. 1 glass bottom $\mu$-Dish (Cellvis) using 1 % low melting point agarose (Promega) in Danieau´s solution (58 mM NaCl, 0.7 mM KCl, 0.4 mM MgSO4, 0.6 mM Ca(NO3)2 and 5 mM HEPES). Before agarose polymerization, embryos were oriented with blastomere cells facing to the glass. Bioluminescence imaging was performed with a 40×/1.25-NA silicon oil-immersion objective, a 100 mm tube lens (ASI Imaging) and a Hamamatsu Orca-Fusion camera (C14440-20UP) was used. Z-stack images of the embryo were taken with 2 s exposure time every 5 $\mu$m, resulting in a volume approximately of 155 $\mu$m. To restore the image, we use the *D. melanogaster* with the membrane marker Ecad::GFP data provided by the authors of ref [16]. We followed their training protocol and inferred our zebrafish luminescence data with the best model checkpoint we obtained. After the projection using the neural networks, we recovered most of the lost cellular membrane information allowing us to perform further analysis such as cellular segmentation. To do this, we applied a morphological dilation filter with kernel size of 3 to join the small gaps where the membrane is not fully complete. Next, we perform watershed segmentation using the Fiji plugin MorphoLibJ to get single cell segment distribution[51].

**Denoising light field images** To denoise light field images we collected a bioluminescent dataset using anesthetized transgenic *C. elegans* under 1 $\mu$l of levamisole 2.5 mM and treated with 1 $\mu$l, 20$\mu$M of fluorofurimazine (FFz, Promega) mounted in 1% of agar pads expressing mTurquoise fused to nanolantern at different developmental stages e.g. L3, L4 and young adults. All subsequent data acquisition were done with the same animal and protocol. To diversify the input distribution of the model, we acquired images with different camera exposure times e.g. 200 ms, 500 ms, 1 sec, 5 sec and 10 sec sequentially, using the same field of view and pairing each image condition to the image with the highest exposure time we collected. For each position and condition we acquired a stack to collect the information of how the light changes through the lenslets over different z positions. For the 3D stacks we took 3 planes above and below the current position with a step size of 1 $\mu$, thereby disentangling the inference model quality from the z position of the bioluminescent sample. For imaging, a bioluminescent light field setup described above was used. We collected 2034 paired of images with a size of 934 $\times$ 976, resulting in $\sim$ 7 GB. We employed the photon number resolving mode to decrease the readout noise for both, training and testing datasets. Next, we employed linear transformations to regularize and prevent overfitting. We used the CARE 2D denoising training topology, extracting 508500 random subimages with a size of 128 $\times$ 128 and a batch size of 64. The training took 15 hours and 32 min in a workstation equipped with Intel(R) Xeon(R) Gold 6248R CPU, 128 Gb of RAM and Nvidia Quadro RTX8000 48GB. All subsequent light field denoising and reconstruction training were done on the same setup. We evaluated the denoised images by calculating the SSIM and RSE metrics in unseen long exposure bioluminescent images (Supplementary Fig 5).

##### 1.0.4 Bioluminescence lightfield reconstruction

For bioluminescence light field reconstruction we use the neural network VCD-Net[37]. VCD-Net is a based U-Net topology composed by a encoder-decoder sampling. The views extracted from the light field images (see Figure 3) are transformed from perspectives to channels intrinsically done by the computation of the convolutional layers. Same as in the original U-Net architecture skip connection are defined between the encoder and decoder to preserve unprocessed information for better fitting. The network gradually transform the extracted views from the original light field image into the conventional 3D stack. The number of filters or dimensionality of the output space in the last convolutional layer is set to match the desired amount of planes for the 3D reconstruction.

**3D reconstruction with synthetic light field data** For the data acquisition for the reconstruction of high quality 3D data from synthetic light field images, we collected high quality 3D stacks, an epifluorescence microscope with 40x$\times$/1.1-NA (numerical aperture) water-immersion objective and 488 nm excitation wavelength were used. We captured 26 different 3D stacks (of size 1024 $\times$ 1024 $\times$ 31). Before the generation of synthetic data, we performed linear transformations, e.g. flipping, rotating and inverting the z axis to augmentate the dataset. Subsequently, we compute the simulated PSF through the microlens-array using [52] to perform a light field projection to the acquired 3D stacks using [37]. This process generates the synthetic light field images that correspond to the input in our neural network. Like the CARE pipeline, the training was done by gradually varying the model coefficients in order to minimize the loss function. For training we extracted 7518 pairs patches of size 176 $\times$ 176 $\times$ 31 pixels and 176 $\times$ 176 pixels for the 3D target and input respectively. We trained for 100 epochs which represented a cost of 7.5 h in a single graphical processing unit. To evaluate the model reconstruction and demonstrate that the relative intensities are preserved, we calculated the SSIM and RSE maps against unseen fluorescence stacks. Furthermore, we compare the intensity profile on one edge of the worm to compare the intensity behaviour between the light field reconstruction and the ground truth (see Supplementary Fig. 4).

**3D reconstruction with experimental light field data via transfer learning** One of the main weakness about training purely with synthetic data is the learning limitation of experimental information that your setup might suffer, such as noise or lens misalignment. Therefore, minor errors introduced in your input might generate artifacts in the prediction. Because light field images and wide field stacks represent optical information in different ways, performing registration between the couple of images in the dataset usually fails. Therefore, the reconstruction of the light field images needs to be made before proceeding to the registration [38]. As we previously stated, the 3D reconstruction of light field images are extremely time consuming. Hence, creating a dataset which consists solely on experimental data where it has enough information to cover the 3D LF reconstruction manifold would represent a gigantic effort. Therefore, we made use of the neural network trained purely with simulated data. For the imaging we used the commercial epifluorescence microscope Leica DMI8 with two available camera ports. In one port, we acquired light field images where we mounted a microlens array of pitch 192 and an effective focal lenght of 3.17 mm (APO-Q-P192-F3.17 (633)) in a CMOS board level camera C11440-62U. The second camera port was dedicated to acquire the corresponding stacks with a CMOS ORCA-Fusion C14440-20UP. An objective 40x/1.1 water immersion lens (Leica) and a 488 nm excitation were employed. We used four different exposure time imaging conditions, e.g. 1 ms, 10 ms, 50 ms and 100 ms for both, stacks and light field images. The same field of view was imaged using light field and epifluorescence senquentially. All conditions were paired to the highest exposure time as GT. We first predicted the experimental light field images to get a rough reconstruction of the structure and then apply an image registration with their corresponding experimental stack. By applying the same transformation to the experiment light field image, the matching training pairs were obtained. In that way, we were able to generate an experimental training dataset composed of $\sim$ 11 GB in matter of hours. We then loaded the weights of the pretrained network with synthetic images and trained again with the well aligned images using the same hyperparameters as we discussed in the previous section. To evaluate the training, we once again calculated the SSIM and RSE metrics against a fluorescent target as our GT. We also compared the reconstruction intensity profile against the GT (Figure S4).

**Bioluminescence light field inference** The bioluminescence light field inferences were made by rectifying the light field image and applying both pipelines consecutively. In other words, aiming to obtain the 3D reconstruction of the bioluminescence scene, the bioluminescence light field image needs to be denoised, rectified and reconstructed respectively. First, the trained 2D denoising CARE pipeline was applied on the raw bioluminescence image to improve the quality and recover lost information due to the noise. Then, we used the open source program LFDisplay [33] to calibrate and locate the microlens array. Next, the rectification process was done in the software provided by [52] consisting in a resampling operation to contain the $11 \times 11$ angular light field views per lenslet.

Finally, the VCD-Net trained with experimental data was used to generate the 3D bioluminescent scene.

#### 1.0.5 Normalization and quantification of errors

**Image normalization** Before training and predicting an image, it is important to normalize it to have a comparable range of intensities. We used the widely used percentile-base normalization [16]

$$N(u; p_{low}, p_{high}) = \frac{u - perc(u, p_{low})}{perc(u, p_{high}) - perc(u, p_{low})}, \tag{1}$$

where $perc(u, p)$ is the *p-th* percentile of all pixel values of $u$. This normalization is done before feeding the images to the network either for training or inference.

**Image quality evaluation** A common image quality metric to compare two images it's *mean squared error (MSE)* which measures the average of the squares of the errors

$$MSE = \frac{1}{n}\sum_{i=1}^{n}(y_i - \tilde{y}_i)^2, \tag{2}$$

however, this metric can't be used without normalize the images since the predicted image $\tilde{y}$, differ considerably compared to the ground truth image $y$. To overcome this, we use the *normalized mean squared error* NRMSE, (range 0-1, lower is better), an image quality assessment defined by [16]. For this, we applied equation 1 and parameterized the prediction with $\gamma(\tilde{y}) = \alpha\tilde{y} + \beta$ to scale the restored image and find the MSE metric where is minimal.

$$NRMSE(y, \tilde{y}) = \sqrt{(MSE(\gamma(\tilde{y}), N(y, 0.1, 99.9)))} \tag{3}$$

where

$$\alpha, \beta = \arg\min_{\alpha', \beta'} MSE(N(y, 0.1, 99.9), \alpha' \tilde{y} + \beta'). \quad (4)$$

To compare structural similarity, we use SSIM [53], which is a measure of the similarity between two images. It has a value of 1 if the images are identical and 0 if they are completely different. We applied the same normalization procedure for the SSIM calculation.

### Supplementary Figures

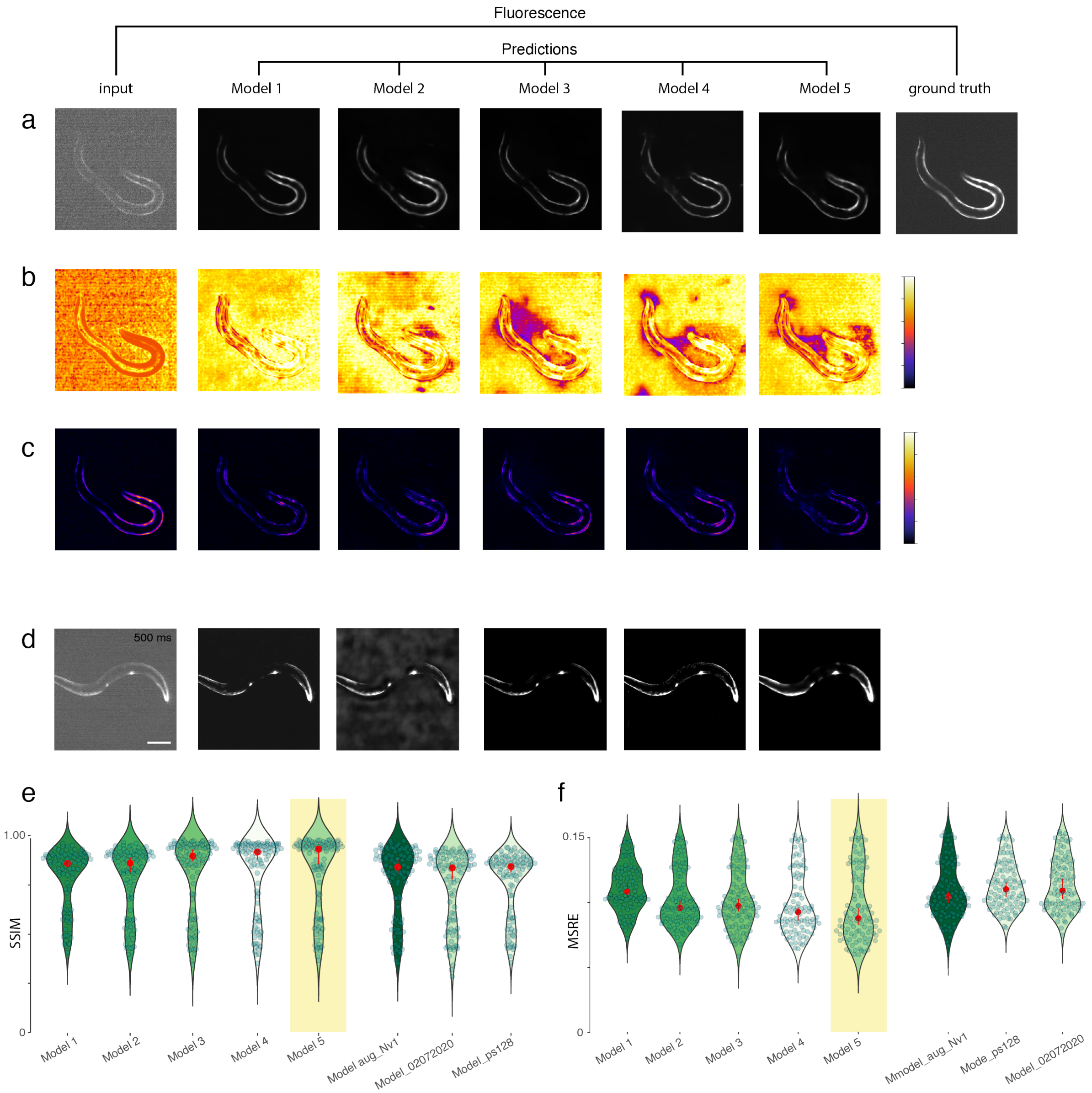


g

| Model | N images | Patches per image | Augmentation | Batch size | Patch size | Percentage Validation |
|---|---|---|---|---|---|---|
| 1 | 2808 | 10 | No | 64 | 64 x 64 | 10 |
| 2 | 2500 | 5 | No | 64 | 128 x 128 | 15 |
| 3 | 25000 | 10 | Yes | 64 | 128 x 128 | 10 |
| 4 | 2500 | 10 | No | 64 | 64 x 64 | 10 |
| 5 | 2500 | 20 | No | 64 | 64 x 64 | 15 |

Supplementary Fig. 1: Benchmarking different models and their qualifiers.
**a,** CARE restorations of the input low SNR image with five best models. **b,** SSIM maps of the input image (left most image) and the different predictions compared to the ground truth, fluorescence image. **c,** RSE maps of the input image (left most image) and the different predictions compared to the ground truth, fluorescence image. **d,** Input bioluminescence image and the inferences of the ground truth for the different models. **e,f,** Summary of the average (e) SSIM and (f) MSRE for all models, tested for 100 different, unseen images. **g** Table of the hyperparameters used to train the different models.

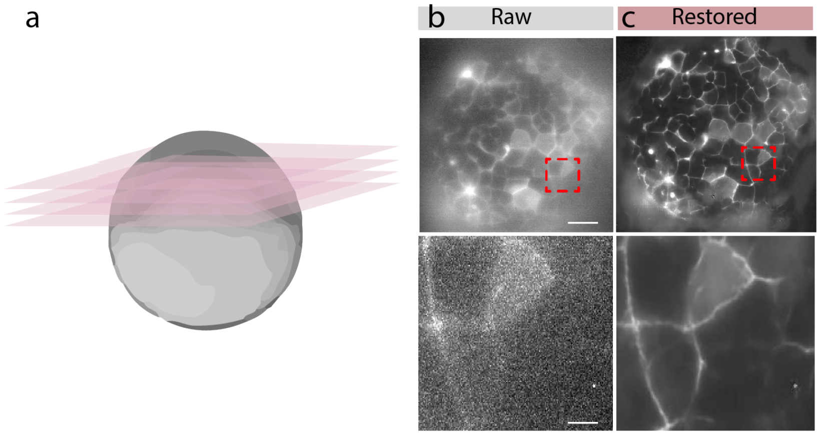


Supplementary Fig. 2: CARE denoising zebrafish embryos improves edge detection
**a,** Schematic of the zebrafish embryo at dome stage indicating the imaging plane of the bioluminescence pictures. **b,** Raw bioluminescence image and high magnification close up.**c,** CARE restoration of the picture in b using a published model for tessellated epithelial monolayers[16]

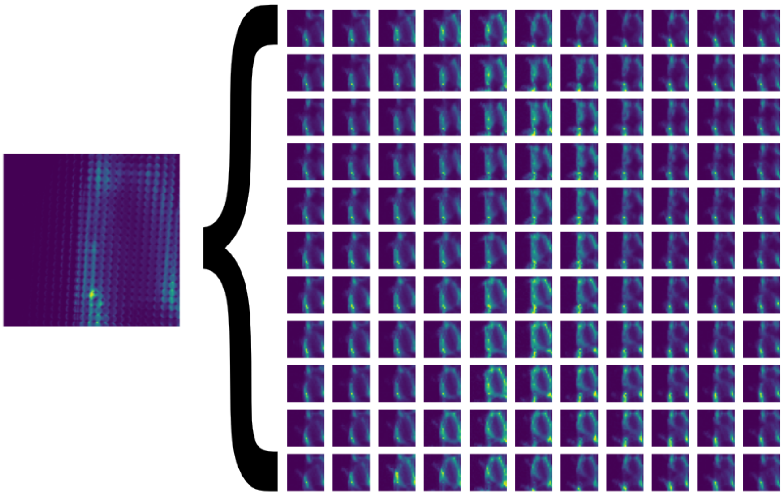

Supplementary Fig. 3: Perspective extraction for lightfield reconstruction
**a,** Extracted perspective views from the bioluminescence picture derived through the microlens array.

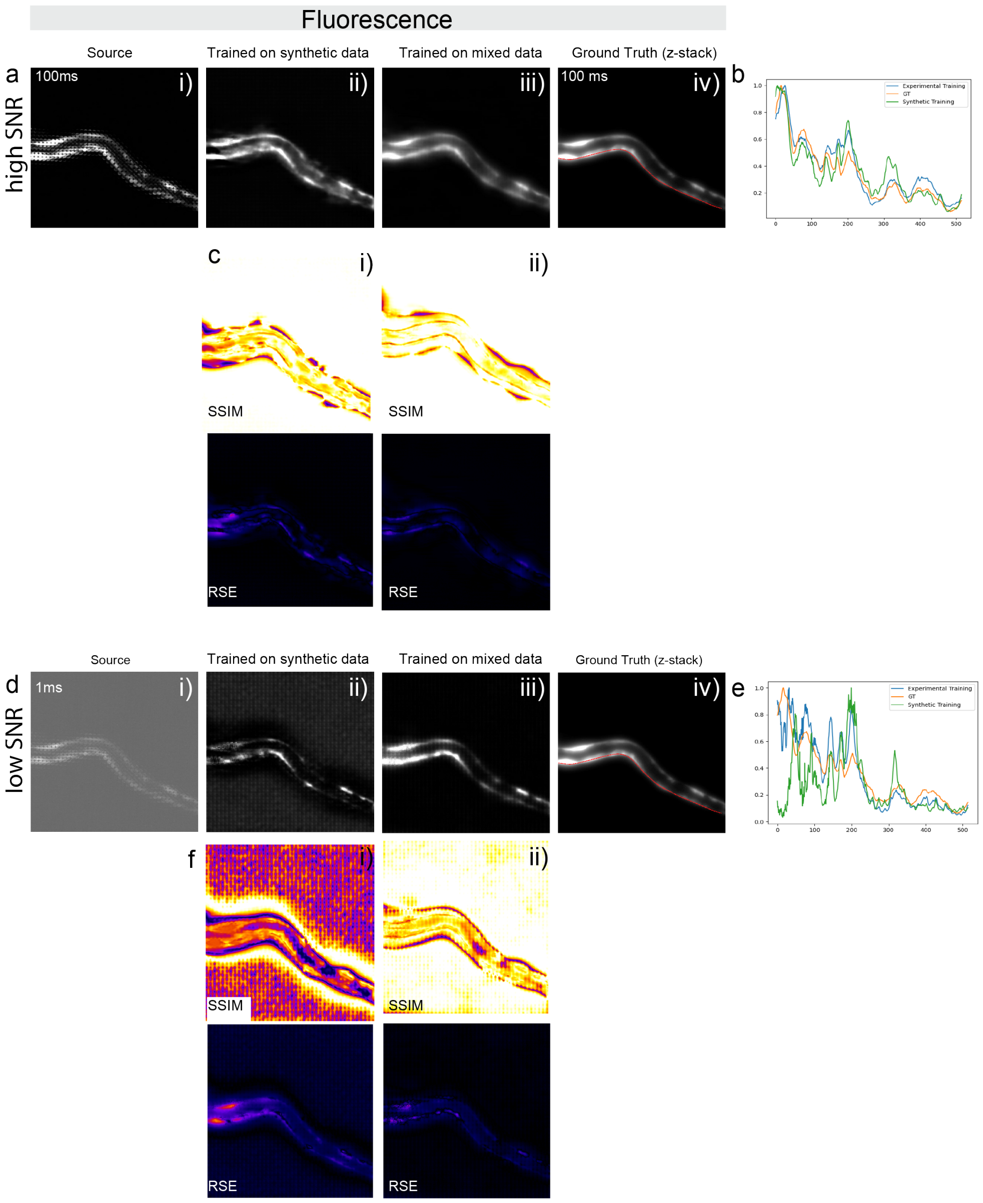


Supplementary Fig. 4: Reconstruction of lightfield images preserves intensity distribution
To test the quality of the training and validity of the model, whether or not the relative input intensities are preserved in the output images, reconstructions of the different models have been tested against the ground truth dataset. **a,** From left to right: i) Input fluorescence lightfield image acquired at 100 ms representing the high SNR image. Reconstruction with a model exclusively trained on ii) synthetic and iii) experimental lightfield data. iv) Maximum intensity projection of a 3D stack acquired with traditional, widefield fluorescence microscopy. **b,** Comparison of the intensity profile for the predicted LF image and the ground truth fluorescence image in the ventral muscles. **c,** SSIM and RSE maps of the i) synthetic and ii) mixed experimental data. **d,** From left to right: i) Input fluorescence image acquired at 1ms representing the low SNR image. Reconstruction with a model exclusively trained on ii) synthetic and iii) experimental lightfield data. iv) Maximum intensity projection of a 3D stack acquired with traditional, widefield fluorescence microscopy. Note, b) iv and d) iv is the same projection from the ground truth image stack. **e,** Comparison of the intensity profile for the predicted LF image and the ground truth fluorescence image in the ventral muscles. **f,** SSIM and RSE maps of the i) synthetic and ii) mixed experimental data.

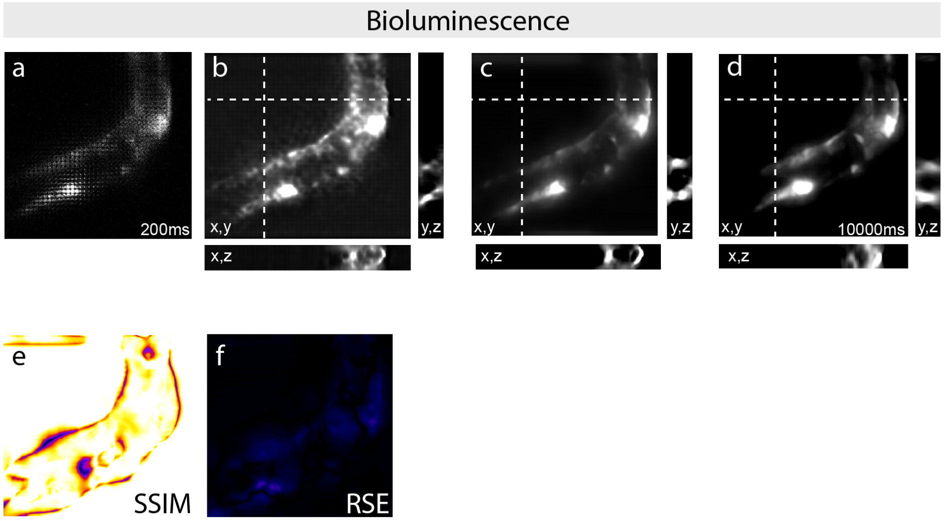


Supplementary Fig. 5: CARE denoising of lightfield images improves 3D reconstructions
**a,** Raw, low SNR light field image of an immobilized bioluminescent animal taken at 200ms exposure time. **b,** Reconstruction without prior denoising. **c,** Reconstruction with a priori CARE denoising. **d,** Reconstruction of a 'ground truth' light field image of the same worm in bioluminescence contrast taken at 10s exposure time. All light field models were trained with mixed (experimental and simulated) data.**e,f** SSIM (e) and RSE (f) of the denoised data tested against the 'ground truth'.
